# Dynamics of genetic circuits in *Pseudomonas protegens*

**DOI:** 10.1101/2024.11.17.623988

**Authors:** Juan Rico, Pablo Japón, Luis Rubio, Ángel Goñi-Moreno

## Abstract

The engineering of genetic circuits to perform predefined computations is central to synthetic biology, enabling living cells with new functionalities applicable across various domains. However, these circuits are often specifically tailored to particular cellular hosts, with *Escherichia coli* being the most popular. Consequently, their intended functions may not translate well to other organisms, limiting their scope. Understanding circuit dynamics in less familiar organisms is crucial, especially for niche-specific applications requiring cellular chassis different from model organisms generally used in synthetic biology. Here, we develop a combined experimental and theoretical pipeline to evaluate the performance of NOT logic circuits, also called inverters, in the soil bacterium *Pseudomonas protegens* Pf-5—a host renowned for its unique environmental functions and a newcomer to genetic circuitry. Inverters were experimentally tested to characterize input-output functionality, and mathematical modelling was used to infer the dynamic principles of circuit modules. The model quantified the individual impacts of key parameters—such as translation efficiency, repressor performance, and promoter activity—on output levels, enabling predictions about inter-host circuit portability. This parameter calibration revealed unique properties of the chassis, including faster transitions between on and off circuit states compared to the synthetic biology workhorse *Pseudomonas putida*. These characteristics may reflect adaptations to the fluctuating conditions of the plant rhizosphere, where this bacteria thrives. As a result, our work provides DNA parts, circuits and mathematical characterizations to establish *P. protegens* Pf-5 as a viable chassis for environmental synthetic biology.

## 1 Introduction

Synthetic biology focuses on the rational design and implementation of predefined computations in living cells [1, 2, 3]. To achieve this goal, the programming of genetic circuits plays a crucial role in sensing inputs and generating outputs according to rules and instructions encoded in DNA sequences [4, 5]. These circuits function as various entities, such as Boolean logic gates [6, 7], counters [8], switches [9], feedback controllers [10], and memories [11], serving as tools to introduce innovative functions into host organisms.

While successful in various applications [12, 13], spanning from biotechnology [14] to bioremediation [15], the programming of biological functions often gives greater emphasis to the software (i.e., the genetic circuit), overlooking the crucial role played by the hardware (i.e., the host cell) [16]. For instance, the optimization of genetic circuits typically relies on testing libraries of DNA parts [17, 18], such as promoters [19] or ribosome binding sites [20], to identify the set of parts that accurately produces the desired phenotypic response. However, it is essential to recognize that the hardware is equally crucial for ensuring the proper performance of the software. For example, certain DNA parts may function effectively in some hosts but poorly in others—the function of genetic circuits is highly context-dependent [21, 22]. This underscores the necessity of tailoring genetic circuits specifically to a particular type of cellular hardware. It could be argued that circuits alone are not sufficient to achieve predefined functionality; rather, the tandem of circuit and host is the actual functional unit. Consequently, what works well in one host often needs to be re-engineered to perform effectively in a different organism [23]. Far from being a disadvantage, this requirement has been leveraged to expand the functional abilities of circuits [21].

Each cellular host possesses specific dynamics that distinguish it from other organisms, ultimately influencing the performance of genetic circuits. While distinctions are more apparent between bacteria and eukaryotic cells, significant mechanistic differences also exist among bacterial species. These distinctions extend beyond important differences in shape [24] or doubling times [25], impacting circuit performance by encompassing variations in resource distribution [26] (e.g., ribosomes) and cellular functions [27] (e.g., transcription vs translation). In synthetic biology, the concept of a chassis [28] refers to a well-characterized host organism that serves as a framework for bioengineering, offering a comprehensive understanding of its impact on the performance of genetic circuits. To date, there is a lack of such well-characterized hosts—a gap that underscores the challenge addressed in this work.

The model bacterium *Escherichia coli* has been pivotal in synthetic biology, benefiting from the extensive knowledge accumulated about this chassis since the early days of molecular biology [29]. However, extending the applications of genetic circuits beyond the laboratory requires customising their functions to suit niche-specific cellular hosts. For instance, there are cells with naturally evolved traits ideal for targeting sustainability applications [30]. Examples include the engineering of cells to linking greenhouse gas sequestration with the bioproduction of valuable compounds using *Clostridium autoethanogenum* and *Cupriavidus necator* [31, 32]. Additionally, rhizosphere-specific applications have been implemented using nitrogen-fixing hosts such as *Azotobacter vinelandii, Stutzerimonas stutzeri*, and *Klebsiella variicola* [33, 34]. Other well-studied cellular chassis for various synthetic biology applications include *Bacillus subtilis* [35], *Pseudomonas putida* [36], *Geobacter sulfurreducens* [37, 38], and non-bacterial organisms like the yeast *Saccharomyces cerevisiae* [39]. In all cases, circuits need to be tailored to their host, not only to adapt to the specific internal workings of the cell but also to leverage this specificity, which may provide advantages in various application domains.

Here, we focus on the soil bacterium *Pseudomonas protegens*, an important environmental host with unique interactions with soil’s biotic (e.g., plants and other microbes) and abiotic (e.g., pollutants) components. For example, this bacterium excels in biocontrol activity against plant pathogens [40, 41]. However, compared to other organisms, the engineering of predefined functions in *P. protegens* has lagged behind. Recent examples, though, highlight its potential as a host for bioengineering endeavours [42, 43]. Despite these advancements, thorough characterization of genetic parts, circuits, context dependencies, and dynamic details is still lacking—an issue that this paper aims to address, unlocking the potential of synthetic biology in this host. To evaluate the context dependency of circuits in *P. protegens*, we performed parallel experiments in the related soil bacterium *P. putida* [44, 45]—a closely related organism with other important environmental traits. By comparing circuit dynamics between the two hosts and developing mathematical models, we were able to accurately predict circuit output before experimental characterization. Although mathematical modelling [46, 47] and computational tools [48, 49] are crucial to genetic circuit design and have become fundamental in synthetic biology [50, 51], such predictive capabilities are still elusive and constitute an overarching challenge in the field [52].

In the following section, we present experimental and theoretical results illustrating the performance of a library of genetic circuits in *P. protegens*. Key dynamic parameters, considered the fingerprint of each circuit, assist in predicting circuit portability between this host and *P. putida*. Altogether, these results contribute to establishing *P. protegens* as a synthetic biology chassis.

## 2 Results and discussion

### Metrics for evaluating genetic circuit performance

To assess the performance of genetic circuits in *Pseudomonas protegens* Pf-5, we used a set of standardized NOT logic gates, or inverters. This benchmarking set is a library of 10 circuits (Figure 1A) that perform a NOT function: if the input regulator is absent (logic 0), the output protein will be expressed at high levels (logic 1), and vice versa. In other words, the output inverts the input.

**Figure 1.**
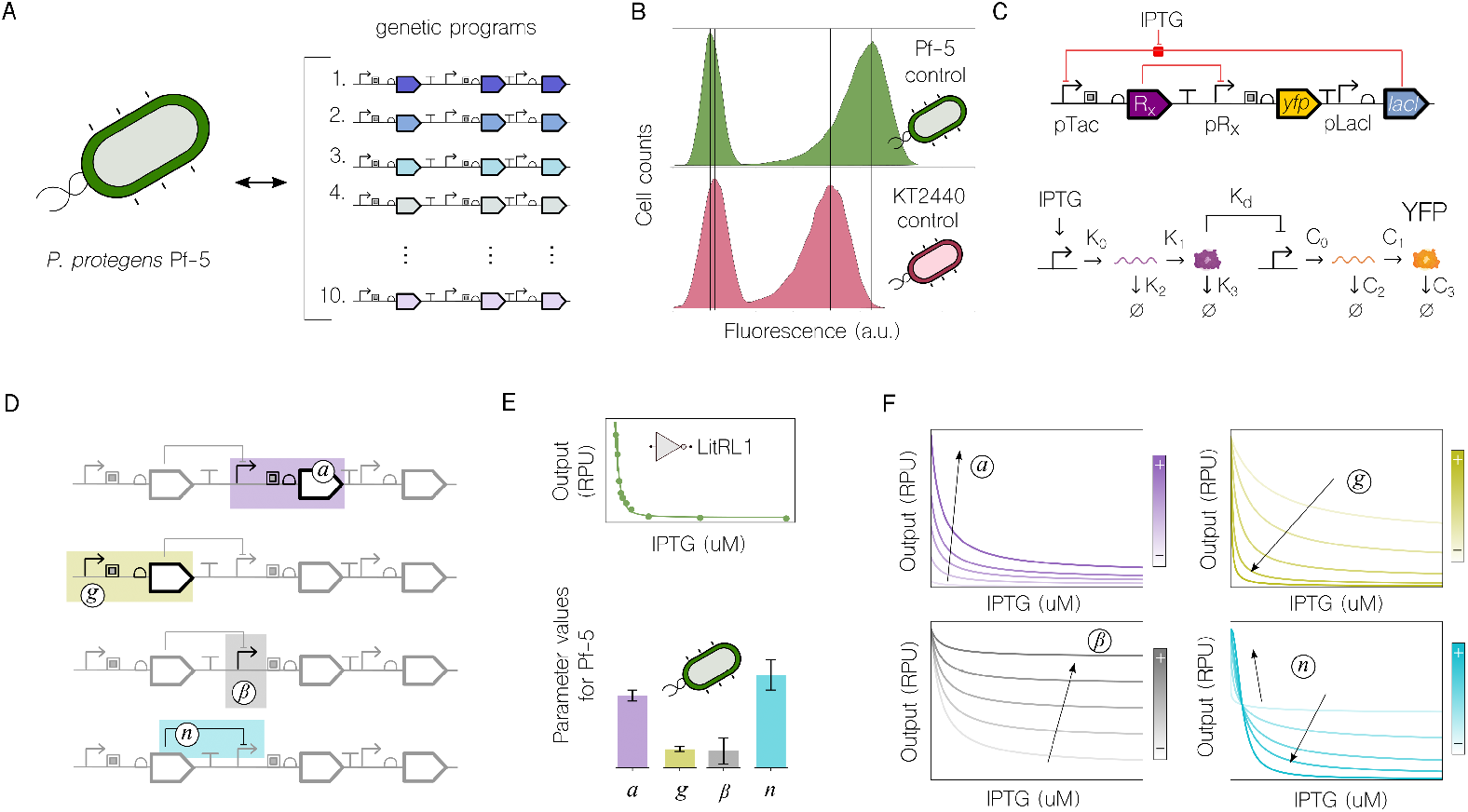
Testing metrics for evaluating genetic circuits performance in different chassis. **A**. A library of 10 genetic NOT logic circuits was tested in *Pseudomonas protegens* Pf-5 at increasing input concentrations, resulting in characterizations of their individual dynamic ranges. **B**. Controls for both the target cell and the reference *Pseudomonas putida* KT2440 in terms of autofluorescence (histogram on the left) and the empty NOT construct with only the reporter (histogram on the right), already exhibit different background performances—which were used to correct circuit output. **C**. The NOT circuits (top) consist of a common architecture shared among all, with a unique set of a repressor and its cognate promoter, denoted here as Rx and pRx, respectively. Circuits were mechanistically modelled (bottom) with the following key rates for parameterizing their performance: K0, K1, K2, K3 and K_*d*_ for the transcription, translation, degradations and dissociation constant of the repressor machinery; and C0, C1, C2, and C3 for the same dynamics involved in YFP expression. Repression was modelled using a Hill equation. **D**. The rates were clustered into 4 testing metrics that can be directly related to the performance of various DNA sequences within the circuits: the reporter expression cassette (*a*), the repressor expression cassette and the Rx-pRx binding dissociation constant (*g*), the leakiness of the central repressible promoter pRx (*β*), and the Rx-pRx binding cooperativity (*n*). **E**. Each NOT logic characterization, resulting in a graph that relates the level of input (IPTG, x-axis) to the level of output (YFP, y-axis), was fitted to the mathematical model, which returns a specific and unique set of testing metrics. **F**. The effect of increasing the value of each of the four metrics on NOT logic output.

Each circuit within the library was independently analysed to measure input-output function. The experimental data were then fitted to the circuit’s mathematical model, and the inverters functionality was compared to the performance of the same testing library in the host *Pseudomonas putida* KT2440. Comparing circuit functions across organisms is challenging because a circuit’s performance is significantly influenced by its host context. To address this, we used measurements of the varying background activities observed in each cell type as controls (Figure 1B). These measurements accounted for both background fluorescence and the signal of an empty NOT gate (i.e., the construct containing only the reporter gene). This activity was used to normalize circuit behaviour against background activity, allowing mathematical modelling to account for both the contextual dependencies impacting the circuit and its intrinsic function. The substantial variance observed in background activity between the two hosts underscores the importance of this step. Although a detailed discussion of the origins of these differences is beyond the scope of this study, we infer that the burden associated with the presence of plasmids may be significantly greater in *P. protegens* compared to *P. putida*. This observation could potentially impact future design strategies.

The structure of the inverter circuits and their mathematical counterparts are depicted in Figure 1C. All synthetic constructs share a common architecture, comprising the reporter gene (*yfp*) used as the output and the repressor-promoter pair (LacI-pTac) that senses input concentration (IPTG). While input and output molecules are the same, the variable component within the library, which defines the unique characteristics of each inverter, is a distinct repressor (Rx) and its corresponding promoter (pRx). The benchmarking set includes seven different repressor-promoter pairs, plus three additional variants for circuits regulated by the QacR or SrpR repressors, making a total of 10 circuits. The mathematical model simulates the NOT function of the gates by considering the mechanistic kinetic rates within each circuit. As depicted in the model scheme, a circuit can be divided into two distinct modules: rates K describe the dynamics of the repressor regulator, while rates C describe those of the reporter. The actual binding and repression of the target promoter pRx is modelled using a Hill equation.

To characterize the gates, models were fitted to the experimental data. However, models based on the full set of rates (K and C) exhibited overfitting, which refers to errors in parameter estimation caused by an excessive number of variables. This occurred because these models were overly complex for the data aimed to match, namely fluorescence measurements. In simpler terms, the models failed to discern the dynamics responsible for variations in performance. Consequently, in order to derive mechanistic insights from the fitting process, we merged all parameters K—along with the Rx-pRx binding dissociation constant—and C into single values, denoted as *g* (repressor cassette) and *a* (reporter cassette), respectively. Additionally, new parameters, denoted as *β* (leakiness) and *n* (Hill coefficient), were introduced to represent the central promoter repressibility (i.e., the leakiness of pRx at maximum repression) and the cooperativity of the repressor binding the output promoter pRx (i.e., the affinity of Rx to bind pRx after other Rx was bound), respectively (Figure 1D). This approach yielded identifiable values that could be fitted to our data and also elucidated mechanistic details inherent to the genetic construct. As a result, for each characterization of a genetic inverter—where input concentration (IPTG) is linked to output expression (YFP) (Figure 1E, top)—the model estimated a specific set of values for the four parameters (*a, g, β, n*) (Figure 1E, bottom). Increasing or decreasing each parameter directly affects inverter functions (Figure 1F), making them suitable for examining experimental results that display differences in curves. For example, increasing parameter *a* shifts inverter curves upwards, particularly at low input (IPTG) values, while high input values result in similar output responses. This occurs because parameter *a* serves as a proxy for the YFP expression machinery; thus, the stronger it is, the more responsive it becomes to low IPTG values (i.e., output 1), while its inhibited state exhibits only a mild increase. Conversely, parameter *g* shifts the curve downwards while maintaining the initial point at low IPTG values relatively constant. The latter happens also in the effect of parameter *β* but shifting the curve upwards. Parameter *n* exhibits a nonlinear impact on the circuit, consistent with what was expected for repression interactions (see Methods - equation 4).

These four metrics effectively described inverter variations and were utilized to analyze the results from the library, as described next.

### Genetic inverters performance in *Pseudomonas protegens* Pf-5

The library of 10 genetic inverters was initially characterized through input-output expression analysis, with Figure 2 showing results normalized against background activity (the RPUs of pTac in contrast to those for each pRx and their experimental variability for *P. protegens* are depicted in Supplementary Figure S1). For each construct, the output (YFP) was measured following induction with increasing concentrations of the input (IPTG). This process generated 10 transfer functions exhibiting distinct differences while maintaining the NOT logic: low IPTG resulted in high YFP and vice versa. Next, experimental observations were used by the mathematical model to determine parameter values for the four performance metrics: *a* (reporter cassette), *g* (repressor cassette), *β* (leakiness), and *n* (Hill coefficient). Error bars in parameter values represent the uncertainty of those parameters. In other words, greater uncertainty indicates that the parameter can vary widely (within the range of the uncertainty) while still fitting the experimental data successfully. Conversely, low uncertainty means the model found a specific value for that parameter. This secondary dataset revealed performance alterations that were not easily estimated through visual inspection alone. For example, certain inverter curves in *P. protegens* may appear similar visually but display significant differences in metric values (e.g., LitRL1 vs QacRQ1).

**Figure 2.**
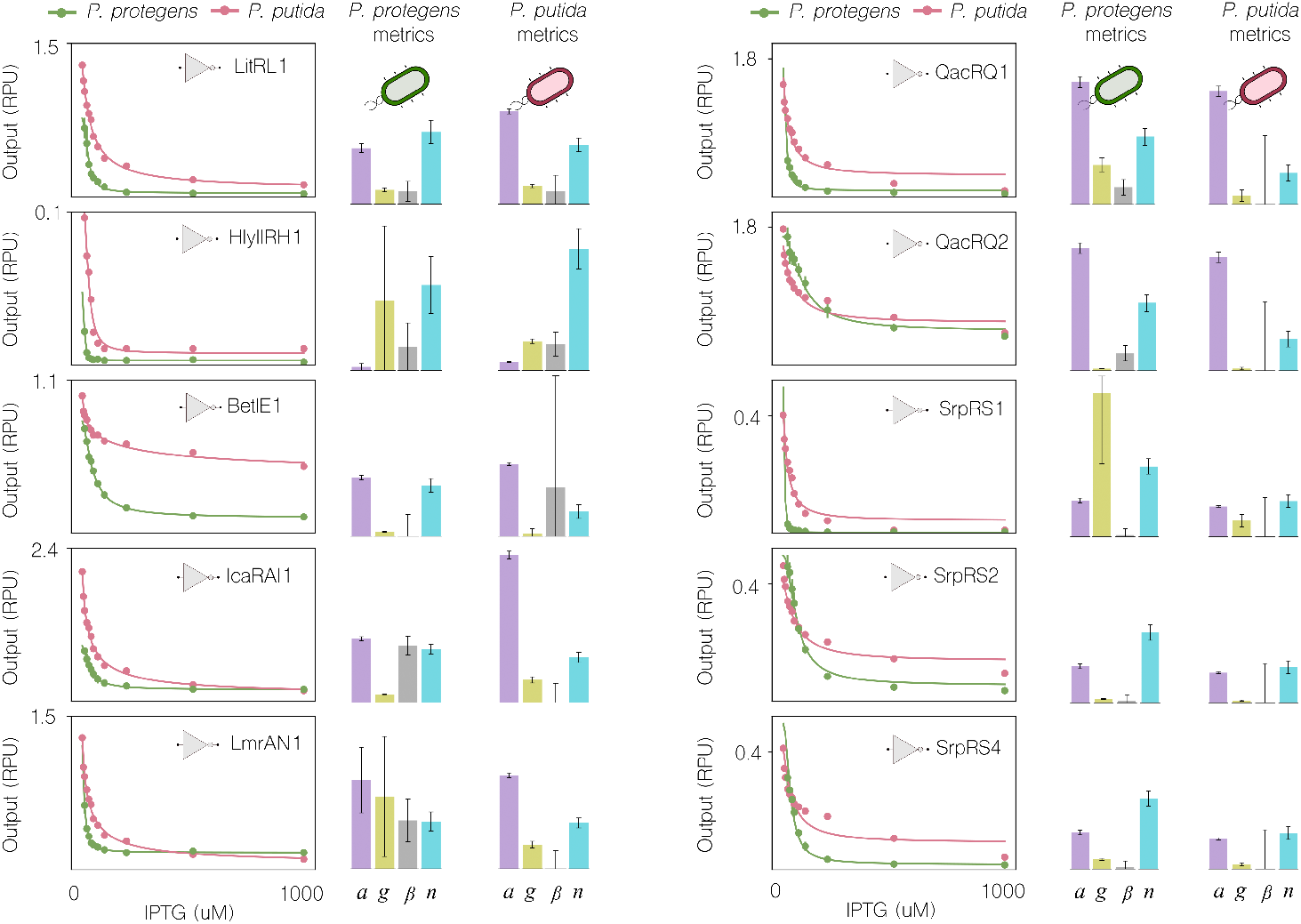
Evaluation of genetic inverters in *P. protegens* Pf-5. Each of the 10 genetic inverters was characterized using input concentrations of IPTG ranging from 0 to 1000 μM. Output fluorescence was measured as relative units adjusting the standards to those of *P. protegens*, facilitating direct comparisons. The results were plotted against those of *P. putida*. In all plots, experimental data and model fit are depicted using dots and solid lines, respectively. Additionally, for each inverter, the 4 testing metrics *a, g, β*, and *n* were calculated and presented in bar plots alongside each inverter function. Parameter values except *g* (inverse concentration arbitrary units) are dimensionless using intervals: 0-2 for *a*, 0-2 (×10^2^) for *g*, 0-4 (×10^*−*1^) for *β*, and 0-3 for *n*.

In this study, the four metrics generated from our combined experimental and theoretical pipeline were used as the unique fingerprint of each genetic inverter. We also applied here this mathematical process to fit previous experimental values in *P. putida* [53], ensuring that the same parameters were obtained to facilitate comparisons.

A closer examination of parameter values and their uncertainty reveals changing levels of variability within the modules of each inverter in the library. For instance, the reporter expression activity, captured by parameter *a*, ranges from 0 to 2 (dimensionless parameter values), with circuits covering most values in between—a common behaviour for both *P. protegens* and *P. putida* (Figure 2). Since the RBS and coding sequence are common to all inverters, this suggests that the variability is mainly due to changes in the promoter, thus offering a wide range of activity to meet future application requirements. Even greater variability is shown by parameter *g*, which encompasses repressor expression and Rx-pRx binding dissociation constant. This ranges from 0 to 100 (inverse concentration arbitrary units). However, *g* displays different performance in the two hosts: while *P. putida* consistently shows relatively low values, *P. protegens* covers the entire range from very low (e.g., circuit BetIE1) to very high (e.g., circuit SrpRS1).

The fact that all circuits share a common backbone makes it easier in some cases to interpret parameter values in terms of mechanistic dynamics. For instance, changes in parameter *a* can be fully attributed to the transcription of the output promoter (rate C0 in the model). However, parameter *g*, which represents the repressor expression cassette, captures the dynamics of only one common DNA part: the IPTG-inducible promoter pTac, used in all inverters. Although this promoter initiates transcription, the RBS and coding sequence are unique to each construct. Therefore, mRNA sequences vary from one repressor to another, affecting rate K0 due to differences in length or complexity. Rates K1, K2, and K3 (Figure 1C) may also differ among the inverters because of variations in translation initiation, protein synthesis, and degradation rates of mRNAs and repressors. These changes in the mechanistic processes underlying parameter *g* (including the repressor-promoter dissociation constant, K_*d*_) likely cause the observed variability in *P. protegens* Pf-5. To provide a mechanistic interpretation, the role of the RBS in the dynamics of parameter *g* is explored later in this study.

The values for parameters *β* and *n* are more uniformly distributed across the ten inverters and do not show much variability (Figure 3A). Despite this, these parameters provide crucial insights and allowed us to draw significant conclusions from their values. Parameter *β* captures the repressibility of the output promoter by its cognate repressor, showing slight variations between 0 and 0.2, with the remarkable exception of gate BetIE1. Unlike the other three parameters, which are unbounded (i.e., their values are not limited to any upper bound), *β* must have a value between 0 and 1, as it represents the percentage relationship between the on and off states of the inverter. Therefore, all inverters perform well in that regard, and the difference between the on and off states is clear and nearly as wide as possible. In the exceptional case of BetIE1, the off state (Figure 2) is only as low as half its maximum on value, indicating that this gate is not working as well as the others in *P. putida*—as a result its *β* is highly uncertain.

**Figure 3.**
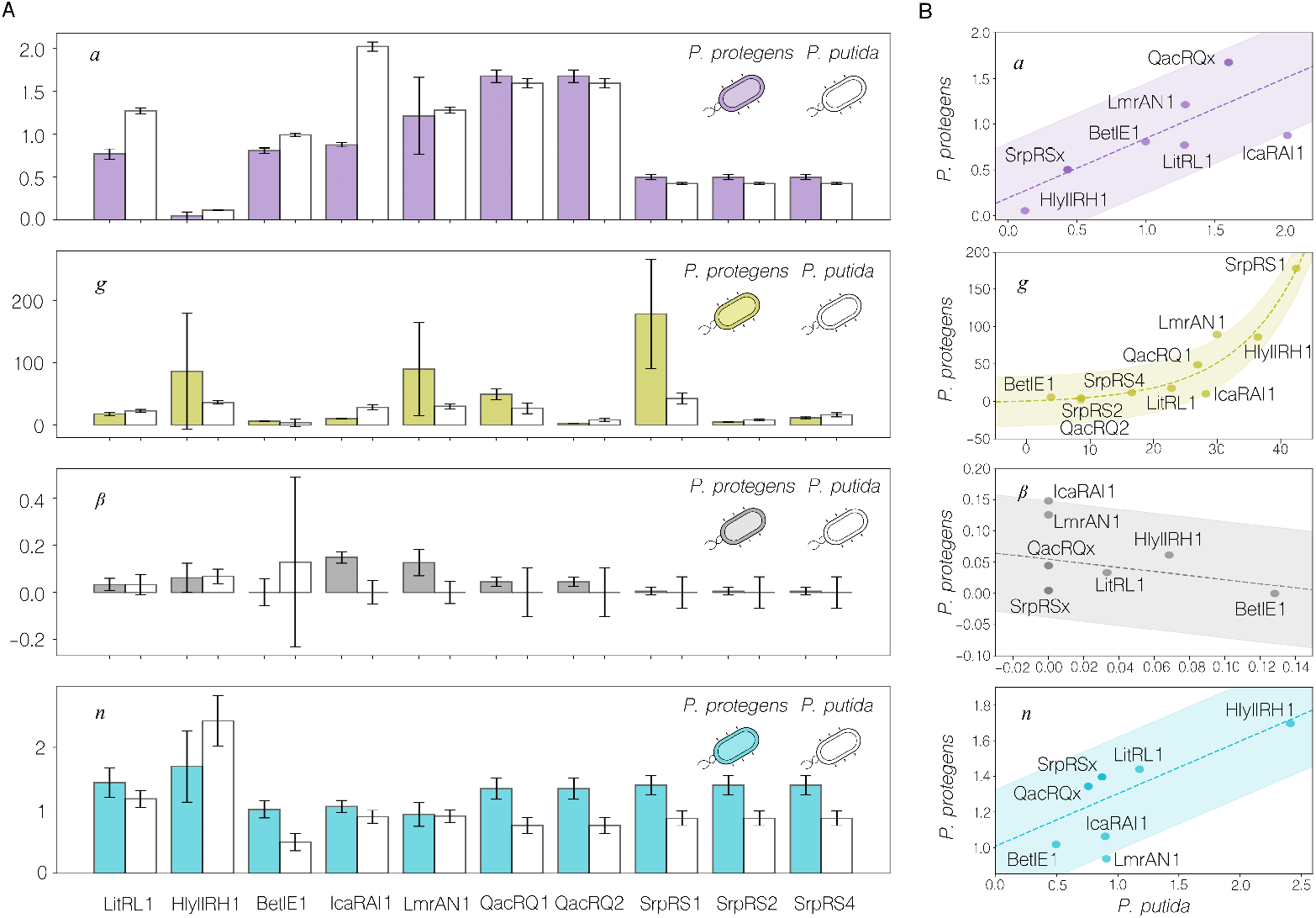
Contextual dependencies of the genetic inverters performance metrics. **A**. Scale of the parameters *a* (purple), *g* (green), *β* (grey) and *n* (blue) extracted from the fitting of the experimental data for each inverter to the mathematical model. The values calculated for each inverter in *Pseudomonas protegens* Pf-5 (coloured bars) are depicted against their equivalents in *Pseudomonas putida* KT2440 (white bars). **B**. Correlations of the inverters performance metrics between *P. putida* KT2440 (x-axis) and *P. protegens* Pf-5 (y-axis). The trajectory and the deviation for each parameter are represented as dashed lines and shadowed areas respectively. Solid dots mark the intersection for each inverter between both hosts. Parameter values except *g* (inverse concentration arbitrary units) in panels A and B are dimensionless.

Parameter *n* showed a clear trend when comparing its values between the two hosts (Figure 3A): it is consistently higher in *P. protegens* Pf-5. Since this parameter captures the cooperativity of the repressor binding to the output promoter within a Hill equation, this higher trend suggests there may be a different regime of nonlinearity between the two organisms. Nonlinear dynamics are crucial for biological systems to run key motifs such as bistability or oscillations, and a consistently higher *n* may provide a library of DNA parts with unique nonlinear trends, aiding the rational design of complex circuits [54].

### Contextual dependencies in *Pseudomonas protegens* Pf-5 compared to *Pseudomonas putida* KT2440

After determining the parameters *a, g, β*, and *n* for each construct, the next question is whether *Pseudomonas protegens* Pf-5 imposes modifications on them that are significantly different from those imposed by the reference strain *Pseudomonas putida* KT2440. While the DNA sequence of the components remains unchanged, their phenotypic response is decisively altered by contextual dependencies. Therefore, from a design perspective, given that DNA sequences can be entirely controlled (mutations and evolutionary dynamics aside), it is desirable to identify the effect of such dependencies.

In order to analyse these dependencies and compare circuit performance metrics between *P. protegens* and *P. putida*, the mathematical model was adjusted to each bacterial host by calibrating the IPTG sensor module shared by all synthetic constructs (LacI-pTac) (Supplementary Figure S2). For each of the four key parameters, circuits displayed different correlation types (Figure 3B): positive linear correlation for parameters *a* and *n*, exponential in the case of *g* and negative linear for *β*. Contextual dependencies are to some extent encoded within these correlations.

For instance, something that may not be obvious looking at Figure 2, where *P. protegens* shows lower initial fluorescence output values in most cases, is that the activity of the output cassettes (parameter *a*) is virtually the same in both hosts. This is inferred by the fact that there is a positive linear correlation in this parameter; besides, values for *P. putida* (x-axis) and *P. protegens* (y-axis) are within the same range. Two additional results strengthen this conclusion, one theoretical one experimental. Firstly, the ratios between rates C0 (output promoter transcription) and K0 (input promoter transcription) also exhibit a similar distribution across inverters and hosts (Supplementary Figure S3). Secondly, the experimentally obtained dynamic ranges (i.e., fold-change between on and off scenarios) are at least as wide in *P. protegens* as in *P. putida* for most inverters (Supplementary Figure S4).

Parameters *β* and *n* are consistently higher in *P. protegens* both concurrently, and linear correlations suggest this trend is stable. This indicates intriguing dynamics: higher *β* values make NOT gates difficult to repress at high input levels, while higher *n* values make NOT gates easy to repress at high input levels (Figure 1F). A potential multi-objective optimization task (beyond the scope of this study) would be to find a sequence that minimizes *β* while maximizing *n* to achieve the best possible NOT function. However, this may not be feasible, as experiments here suggest the host context influence both parameters simultaneously.

However, it is parameter *g* that shows the most significant performance differences due to context effects: exponential correlation and very different range values on the axis. We now take a closer look at this parameter, which accounts for the translation dynamics of the repressor.

### The *P. protegens* Pf-5 context enhances the RBS effect over the repressor expression cassette activity

There are 5 variants within the library of inverters (2 regulated by QacR and by SrpR) which only differ on the RBS sequence of the repressor. Figure 4A sketches the general architecture of these inverters and the mechanistic rates integrating the parameter *g*. The plots of either QacRQ1 and QacRQ2 (Figure 4B) or SrpRS1, SrpRS2 and SrpRS4 (Figure 4C) were merged for both *P. protegens* Pf-5 (top) and *P. putida* KT2440 (bottom) along with the four performance metrics for each host. The visual inspection of the inverter functions show that altering the repressor RBS sequence is enough to modify the input-output relationship—with this effect being clearly more accused in *P. protegens* Pf-5. Moreover, this correlates with the changes attributed by the model to the activity (parameter *g*) of the repressor expression cassette.

**Figure 4.**
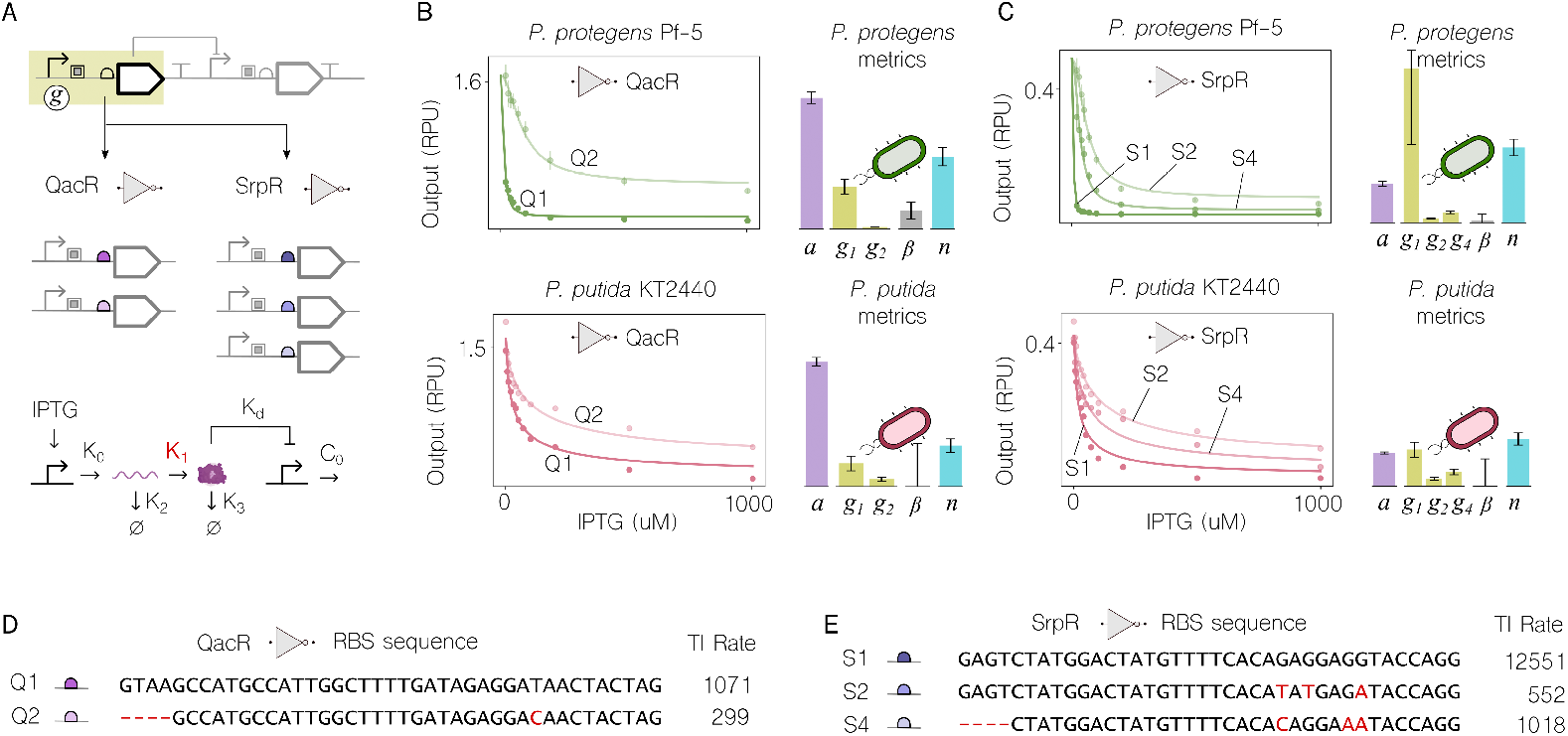
Differential impact of RBS sequence over parameter *g*. **A**. Schematic representation of the inverters architecture highlighting the main genetic elements integrating the repressor expression cassette, thus impacting the parameter *g* (top); draft of the QacR and SrpR repressor expression cassette variants (middle); detail on the mechanistic processes involved in the repressor expression cassette activity (i.e., transcription, translation, mRNA degradation, protein degradation and dissociation constant), and their rates (K0, K1, K2, K3 and K_*d*_) showcasing protein synthesis (K1 – in red) as a key process (bottom). **B**. Experimental data and model fits of the QacR-based inverter variants in *P. protegens* Pf-5 (green) and *P. putida* KT2440 (red) are depicted using dots and solid lines, respectively. The 4 testing metrics *a, g, β*, and *n* were calculated and presented in bar plots - *g*_*1*_ and *g*_*2*_ correspond to Q1 and Q2 RBS variants respectively. **C**. Experimental data and model fits of the SrpR-based inverter variants in *P. protegens* Pf-5 (green) and *P. putida* KT2440 (red) are depicted using dots and solid lines, respectively. The 4 testing metrics *a, g, β*, and *n* were calculated and presented in bar plots - *g*_*1*_, *g*_*2*_ and *g*_*4*_ correspond to S1, S2 and S4 RBS variants respectively. Parameter values except *g* (inverse concentration arbitrary units) in panels B and C are dimensionless using intervals: 0-2 for *a*, 0-2 (×10^2^) for *g*, 0-4 (×10^*−*1^) for *β*, and 0-3 for *n*. **D**. Nucleotide sequences of the RBS variants and their predicted translation initiation rate for the QacR-based inverters. **E**. Nucleotide sequences of the RBS variants and their predicted translation initiation rate for the SrpR-based inverters. Positions with nucleotide substitutions or deletions are highlighted in red in panels D and E.

The predicted translation initiation rates (TIRs) of the mRNA variants for QacR and SrpR were calculated through the De Novo DNA [55, 56, 57] algorithms (Supplementary Table S3). An extract of the relevant information including the name, differential sequence and TIR is also presented in Figure 4D (QacR) and 4E (SrpR)—of note that the calculated TIR was the same regardless of the bacterial host selected. In general terms, if we compare the predicted TIRs with the *g* values calculated through our experimental and theoretical pipeline, both (TIR and *g* values) follow the same trend: the higher the predicted TIR, the higher also the value obtained for *g*. This result supports our model as a tool to evaluate functional modules within genetic circuits while inferring the impact of individual rates (e.g., protein translation) over the modules and the circuits themselves.

Modulation of protein accumulation or circuit performance through RBS modifications is not unconventional. However, here we go a step further by providing a framework to study the effect of such modifications in a context-dependent manner. Our results suggest that there is an intrinsic contextual effect over RBS modifications on phenotypic variance. As a consequence, the cell’s translation machinery can be leveraged to control how RBS modifications influence input-output functions as desired. Indeed, RBS changes produce an estimated 40-fold (S1/S2) increase in the overall repressor expression cassette activity for SrpR when implemented as a *P. protegens* Pf-5 logic circuit, in contrast to the modest (5-fold) phenotypic response observed within the context of *P. putida*.

Nonetheless, this result could be attributed to more than the TIR alone. For instance, translation processivity could favor the RBS effect. We calculated the codon adaptation index (CAI) for the 7 NOT repressors, YFP, and LacI in both bacterial species [58]. The resulting CAI (Supplementary Table S4) is lower in *P. protegens* Pf-5 for all of them. These results raise intriguing questions: Are RBS-related dependencies linked to differences in protein translation across hosts, or are they buffered by other factors like mRNA levels and protein degradation? While these remain unanswered, our findings suggest that combining RBS designs with host context is key to improving genetic circuit portability. A recent study also supports this, showing that while RBS modulation fine-tunes function within a host, considering host context can shift overall circuit performance [59]. Despite some uncertainties about the underlying mechanisms, we can leverage this phenomenon to engineer circuits and even predict their function in different hosts with varying intrinsic properties.

### Prediction of circuit portability between *P. putida* KT2440 and *P. protegens* Pf-5

Portability between bacterial hosts remains a major challenge for the application of synthetic genetic circuits. What functions well in one host often fails in another. The ability to predict a circuit’s performance in a new host would be highly valuable to the field. Here, we developed a predictor tool based on the four parameters *a, g, β*, and *n*.

Figures 5A–C illustrate the process of predicting the function of a circuit tested in *P. putida* KT2440 and applying it to *P. protegens* Pf-5. The LitRL1 gate was selected for prediction (Figure 5A). The parameter values of the remaining nine gates in the library were correlated between *P. putida* and *P. protegens*, providing a setup to interpolate *P. putida* LitRL1 parameter values and predict the parameters for this gate in *P. protegens* (Figure 5B). Importantly, LitRL1 was left out of this correlation process. Leave-One-Out Cross-Validation (LOOCV) [60] is a model evaluation technique where the model is trained on n-1 samples, with one left out as the test set. Figure 5C shows the prediction of how LitRL1 will perform in *P. protegens* (black line) compared against the experimental data of LitRL1 in that host (green dots), showing good agreement between them.We assessed prediction accuracy by comparing predicted and experimental fits, with an R^2^ value above 0.8 for this case (Supplementary Table S5), where 1 represents the best possible outcome.

**Figure 5.**
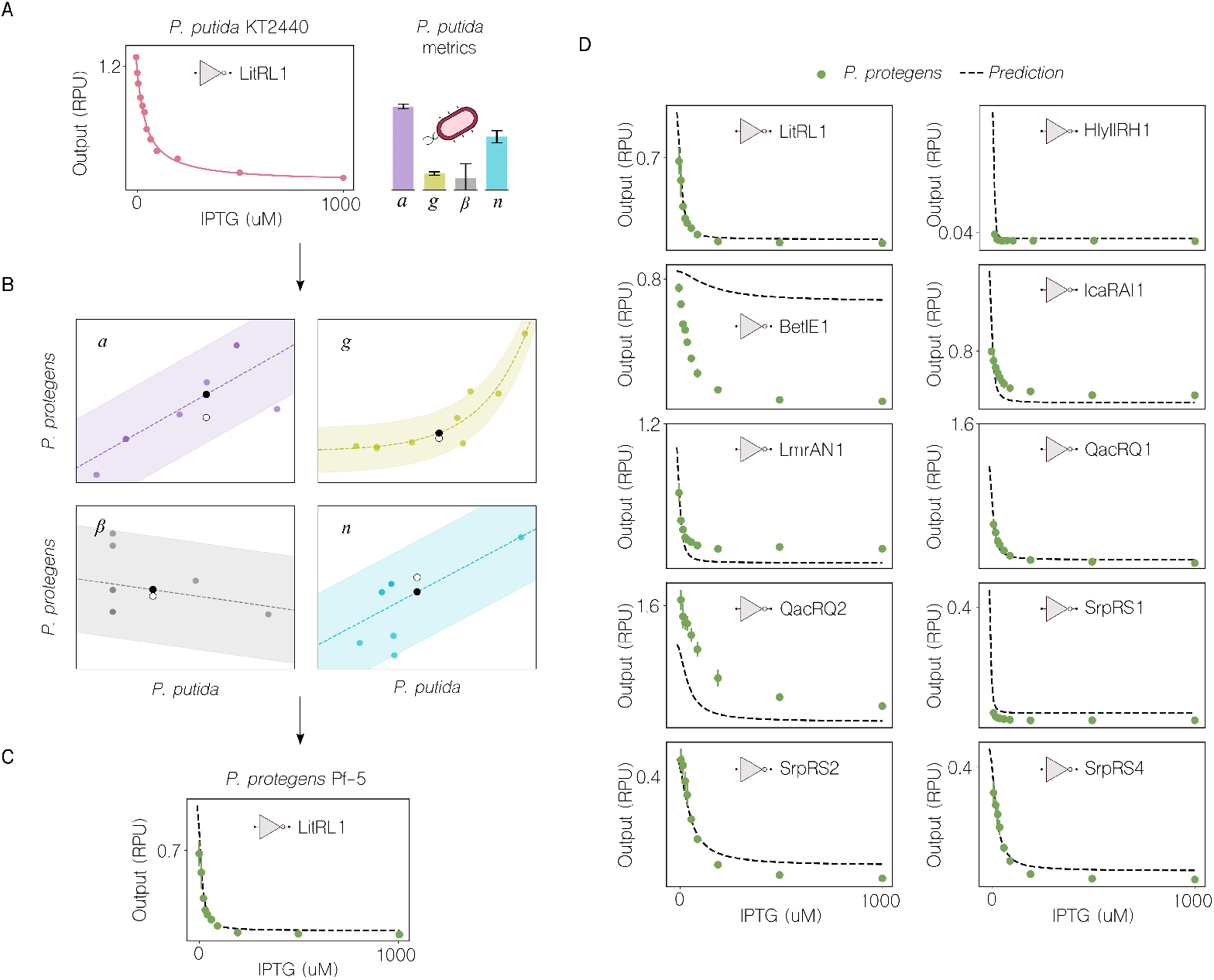
Roadmap for the prediction of circuit portability between hosts. Panels A, B and C represent a visual breakdown of the process from experimental data to inter-host prediction of inverter dynamics. **A**. Experimental data and model fit of the LitRL1 inverter in *P. putida* KT2440 are depicted using dots and solid lines, respectively. Parameter values except *g* (inverse concentration arbitrary units), represented as bars are dimensionless using intervals: 0-2 for *a*, 0-2 (×10^2^) for *g*, 0-4 (×10^*−*1^) for *β*, and 0-3 for *n*. **B**. Interpolation of the parameters (*a, g, β* and *n*) from the LitRL1 inverter in *P. putida* KT2440 into the correlations generated with the other nine gates (all but LitRL1); the predicted and the experimentally-assessed intersections for each parameter between *P. putida* KT2440 (x-axis) and *P. protegens* Pf-5 (y-axis) are represented as black and white dots, respectively. **C**. The predicted LitRL1 inverter dynamics in *P. protegens* Pf-5 is plotted along with the experimental data as a black dashed line and green dots, respectively. **D**. Predicted inverter dynamics and experimental data represented as black dashed lines and green dots for each of the 10 NOT logic circuits included in this study using the correlations of the other nine.

This process was repeated for all 10 gates in the library, completing the LOOCV technique to thoroughly assess predictability (full results in Supplementary Figure S5). As shown in Figure 5D, four gates were exceptionally well predicted, with confidence metrics (Supplementary Table S5) exceeding 0.8 (out of 1) for LitRL1, QacRQ1, SrpRS2, and SrpRS4. SrpRS1 had an intermediate score of 0.69, while the remaining gates showed low to very low predictability. These results mark a significant step forward in predicting genetic circuit performance in P. protegens Pf-5.

Besides, a simulation of 1000 inverters with parameters within the ranges of the benchmarking library utilized in this study was run to confirm or reject the model-based predictions. The probability of achieving predictions with confidence metrics within the range of those obtained in this study was below 0.05 after excluding the HlyIIRH1 and BetIE1 outliers (Supplementary Figure S6). This result validates our roadmap towards the prediction of circuits portability to novel synthetic biology chassis.

## 3 Conclusion

Studying non-model species is crucial for applying synthetic biology tools and engineering them effectively. In this study, we characterized a library of genetic circuits (NOT logic gates) in the soil bacterium *Pseudomonas protegens* Pf-5, examining their dynamics, host-circuit interactions, and developing parameterized mathematical models specific to this organism. Additionally, we developed a method to predict the performance of circuits previously tested in *Pseudomonas putida* KT2440 within this new chassis.

The mathematical model was able to quantify the activity of key circuit components: the reporter cassette (for readout), repressor cassette (implementing the NOT logic), promoter leakiness, and the Hill coefficient (for repressor interactions). Fluorescence measurements for each gate were fitted to these parameters, yielding unique values for each construct. Analysis of these values led to novel mechanistic insights about *P. protegens* Pf-5 that are crucial for the success of bioengineering efforts. Our results suggest that the LacI-pTac inducible system is highly sensitive to very low IPTG concentrations in this host. Given that low-dose systems are advantageous but challenging to engineer, as recently in *E. coli* [61], this trait positions *P. protegens* as a promising chassis to this end. Furthermore, RBS sequences with high TIR efficiently initiate translation in this host, which, along with the system’s sensitivity to low IPTG levels, may explain the narrower transition phase compared to the synthetic biology chassis *P. putida* KT2440. This characteristic is beneficial for applications requiring a digital input-output response [62, 63]. The consistently high Hill coefficients across all circuit models suggest a strong non-linear behaviour in this bacterium, which is key dynamical feature to consider when designing and implementing genetic circuits. All this information—and more that is described in the Results section—aids engineering and enhances predictability in this unconventional chassis.

Bioengineering applications targeting ecosystems need environmental microbes as workhorses. We advocate for the synthetic biology potential of *Pseudomonas protegens* Pf-5 to engineer beneficial traits in soil environments.

## 4 Materials and Methods

### Strains, media and general culture conditions

The bacterial strains used in this study are detailed in the Supplementary Tables S1 and S2. As a general overview, *Escherichia coli* CC118*λpir* with inverters from the pSEVA221 library previously used by Tas *et al* [21] were used as donor strains in triparental matings with *P. protegens* Pf-5 as recipient strain in a similar manner to that previously reported [64]. In brief, *E. coli* CC118*λpir* carrying either the control plasmids pSEVA221:1201, pSEVA221:1717, pSEVA221:1818 or the pSEVA221:Inverters (Supplementary Table S1) were mixed with the helper strain *E. coli* HB101/pRK600 and the recipient strain *P. protegens* Pf-5 in a 1:1:1 ratio based on their OD_600_. All cells were washed with 10 mM MgSO_4_ prior to mixing to reduce the amount of antibiotics carried from the culture medium. The cell suspensions were then centrifuged for 5 minutes at 5000g, resuspended in 10–20 μL of 10 mM MgSO_4_ and a single drop was laid in the center of a Luria-Bertani agar plate. Plates were incubated at 30°C for 20-22 h and the whole cell-patch was resuspended in 1 ml of 10 mM MgSO_4_. Undiluted cell suspensions or up to 10^*−*3^ dilution were plated in M9-citrate agar plus kanamycin 50 μg/ml. Colonies were clearly visible after 24-48 h of incubation at 30ºC. Whole Plasmid Sequencing was performed by Plasmidsaurus using Oxford Nanopore Technology with custom analysis and annotation.

### Phenotypic characterization

#### Induction assays for flow cytometry assessment of gate performance

Bacterial strains of *P. protegens* Pf-5 (Supplementary Table S2) were grown in M9 medium supplemented with citrate as previously described [53]. In summary, saturated overnight cultures were diluted fold and 3 μl were inoculated into the wells of a microtiter plate to a final volume of 200 μl per well, with IPTG concentrations ranging 0 to 1000 μM and incubated at 30°C with shaking for 24 h. Cultures were then kept on ice for the rest of the procedure. YFP fluorescence distribution of each sample was measured with a Miltenyi Biotec MACSQuant VYB flow cytometer at channel B1 with an excitation of 488 nm and emission of 525/50 nm. For each sample 30 thousand events were collected with singlet gating. Calibration was done by using MACSQuant Calibration Beads.

### Data analysis

#### Flow cytometry

The medians for the YFP (B1 channel) fluorescence distribution for each *P. protegens* Pf-5 listed at Supplementary Table S2 were calculated using the MACSQuantify software V2.13.3. The singlet gating dataset for each IPTG concentration (ranging from 0 to 1000 μM) was analyzed.

#### Assessment of the relative promoter units (RPU)

The medians obtained from the flow cytometry data analysis were included in the following formula to assess the relative promoter units (RPU):

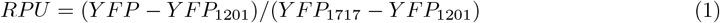

#### Control Model

A typical Hill function where IPTG induces the production of YFP has been used for the control gate (1818).

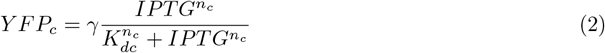

The parameters of the control model are:

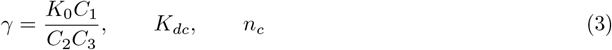

where *γ* is the maximum theoretical *Y FP*_*c*_, *K*_*dc*_ is the dissociation constant and *n*_*c*_ is the Hill coefficient. *K*_*dc*_ and *n*_*c*_ values are used for fitting the NOT gates.

#### Model

NOT gates are modelled by:

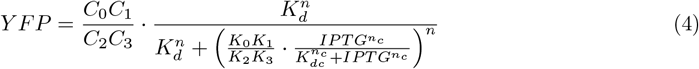

Given our data, we cannot differentiate between some parameters such as *C*_0_ and *C*_1_, that is why we group certain parameters into other new parameters, so that the fit is performed concisely and the result is reliable, avoiding overfitting. Thus:

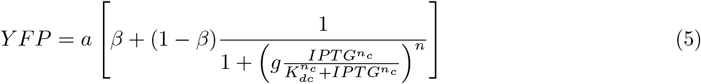

*β* has been added as a measure of gate leakage (*Y FP* (*IPTG >>*)).

the parameters of the model used are:

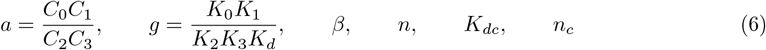

where *a* and *g* group undecidable parameters, *β* refers to the leakage percentage (from 0 to 1), *n* is the Hill coefficient, and *K*_*dc*_ and *n*_*c*_ are the Hill parameters of the control gate. *K*_*dc*_ and *n*_*c*_ are fixed parameters for each host given by the control model (Supplementary Figure S2).

## Acknowledgments

This work was supported by the the ECCO (ERC-2021-COG-101044360) Contract of the EU and grants MULTI-SYSBIO (PID2020-117205GA-I00), BIOELECTRIC (CNS2022-135951), and MULTISYNBIO (PID2023-152470NB-I00) funded by MICIU/AEI/ 10.13039/501100011033.

