## Supplementary material for "Dynamics of genetic circuits in *Pseudomonas protegens*": Supp Info

Table 1: **Donor strains to conjugate the library into *Pseudomonas protegens* Pf-5**.

| **Name** | **Strain** | **Plasmid** | **Ab** | **Function** | **Source** |
|---|---|---|---|---|---|
| **Tas65** | *Escherichia coli* CC118$\lambda pir$ | pSEVA221::1201 | Km | Autofluorescence | [1] |
| **Tas68** | *Escherichia coli* CC118$\lambda pir$ | pSEVA221::1717 | Km | Standardization | [1] |
| **Tas317** | *Escherichia coli* CC118$\lambda pir$ | pSEVA221::1818 | Km | Promoter activity | [1] |
| **Tas3** | *Escherichia coli* CC118$\lambda pir$ | pSEVA221::BetIE1 | Km | Inverter | [1] |
| **Tas7** | *Escherichia coli* CC118$\lambda pir$ | pSEVA221::HlyIIRH1 | Km | Inverter | [1] |
| **Tas8** | *Escherichia coli* CC118$\lambda pir$ | pSEVA221::IcaRAI1 | Km | Inverter | [1] |
| **Tas9** | *Escherichia coli* CC118$\lambda pir$ | pSEVA221::LitRL1 | Km | Inverter | [1] |
| **Tas10** | *Escherichia coli* CC118$\lambda pir$ | pSEVA221::LmrAN1 | Km | Inverter | [1] |
| **Tas15** | *Escherichia coli* CC118$\lambda pir$ | pSEVA221::QacRQ1 | Km | Inverter | [1] |
| **Tas16** | *Escherichia coli* CC118$\lambda pir$ | pSEVA221::QacRQ2 | Km | Inverter | [1] |
| **Tas17** | *Escherichia coli* CC118$\lambda pir$ | pSEVA221::SrpRS1 | Km | Inverter | [1] |
| **Tas18** | *Escherichia coli* CC118$\lambda pir$ | pSEVA221::SrpRS2 | Km | Inverter | [1] |
| **Tas20** | *Escherichia coli* CC118$\lambda pir$ | pSEVA221::SrpRS4 | Km | Inverter | [1] |

Table 2: **Strains and plasmids used for the characterization of inverters.**

| Name | Strain | Plasmid | Ab | Function | Source |
|---|---|---|---|---|---|
| **Tas257** | *Pseudomonas putida* KT2440 | pSEVA221::1201 | Km | Autofluorescence | [1, 2] |
| **Tas260** | *Pseudomonas putida* KT2440 | pSEVA221::1717 | Km | Standardization | [1, 2] |
| **Tas266** | *Pseudomonas putida* KT2440 | pSEVA221::1818 | Km | Promoter activity | [1, 2] |
| **Tas195** | *Pseudomonas putida* KT2440 | pSEVA221::BetIE1 | Km | Inverter | [1, 2] |
| **Tas199** | *Pseudomonas putida* KT2440 | pSEVA221::HlyIIRH1 | Km | Inverter | [1, 2] |
| **Tas200** | *Pseudomonas putida* KT2440 | pSEVA221::IcaRAI1 | Km | Inverter | [1, 2] |
| **Tas201** | *Pseudomonas putida* KT2440 | pSEVA221::LitRL1 | Km | Inverter | [1, 2] |
| **Tas202** | *Pseudomonas putida* KT2440 | pSEVA221::LmrAN1 | Km | Inverter | [1, 2] |
| **Tas207** | *Pseudomonas putida* KT2440 | pSEVA221::QacRQ1 | Km | Inverter | [1, 2] |
| **Tas208** | *Pseudomonas putida* KT2440 | pSEVA221::QacRQ2 | Km | Inverter | [1, 2] |
| **Tas209** | *Pseudomonas putida* KT2440 | pSEVA221::SrpRS1 | Km | Inverter | [1, 2] |
| **Tas210** | *Pseudomonas putida* KT2440 | pSEVA221::SrpRS2 | Km | Inverter | [1, 2] |
| **Tas212** | *Pseudomonas putida* KT2440 | pSEVA221::SrpRS4 | Km | Inverter | [1, 2] |
| **JR230** | *Pseudomonas protegens* Pf-5 | pSEVA221::1201 | Km | Autofluorescence | This study |
| **JR231** | *Pseudomonas protegens* Pf-5 | pSEVA221::1717 | Km | Standardization | This study |
| **JR232** | *Pseudomonas protegens* Pf-5 | pSEVA221::1818 | Km | Promoter activity | This study |
| **JR243** | *Pseudomonas protegens* Pf-5 | pSEVA221::BetIE1 | Km | Inverter | This study |
| **JR239** | *Pseudomonas protegens* Pf-5 | pSEVA221::HlyIIRH1 | Km | Inverter | This study |
| **JR247** | *Pseudomonas protegens* Pf-5 | pSEVA221::IcaRAI1 | Km | Inverter | This study |
| **JR234** | *Pseudomonas protegens* Pf-5 | pSEVA221::LitRL1 | Km | Inverter | This study |
| **JR248** | *Pseudomonas protegens* Pf-5 | pSEVA221::LmrAN1 | Km | Inverter | This study |
| **JR252** | *Pseudomonas protegens* Pf-5 | pSEVA221::QacRQ1 | Km | Inverter | This study |
| **JR236** | *Pseudomonas protegens* Pf-5 | pSEVA221::QacRQ2 | Km | Inverter | This study |
| **JR237** | *Pseudomonas protegens* Pf-5 | pSEVA221::SrpRS1 | Km | Inverter | This study |
| **JR240** | *Pseudomonas protegens* Pf-5 | pSEVA221::SrpRS2 | Km | Inverter | This study |
| **JR242** | *Pseudomonas protegens* Pf-5 | pSEVA221::SrpRS4 | Km | Inverter | This study |

Table 3: **De Novo DNA prediction of translation initiation.** [3, 4, 5]

| | **QacRQ1** | **QacRQ2** | **SrpRS1** | **SrpRS2** | **SrpRS4** |
|---|---|---|---|---|---|
| **Initial State Free Energy** (kcal/mol) | -23.52 | -25.81 | -25.24 | -22.63 | -24.30 |
| **Final State Free Energy** (kcal/mol) | -23.21 | -22.66 | -30.40 | -20.85 | -23.87 |
| **Total Free Energy Change** (kcal/mol) | 0.31 | 3.15 | -5.16 | 1.78 | 0.43 |
| **Translation Rate Prediction** (a.u.) | 1070.98 | 298.68 | 12551.31 | 552.40 | 1018.11 |
| **Shine-Dalgarno Interaction** (kcal/mol) | -5.43 | -5.43 | -12.02 | -2.47 | -4.96 |

Table 4: **Codon Adaptation Index.** $^{a}$percentage. [6]

| Name | Length (bp) | CAI-KT2440 | CAI-Pf-5 | $^{a}$GC | $^{a}$GC1 | $^{a}$GC2 | $^{a}$GC3 | Nc |
|---|---|---|---|---|---|---|---|---|
| **BetI** | 585 | 0.518 | 0.416 | 51.5 | 66.7 | 45.1 | 42.6 | 22.4 |
| **HlyIIR** | 636 | 0.613 | 0.541 | 40.9 | 46.7 | 27.8 | 48.1 | 29.3 |
| **IcaRA** | 561 | 0.650 | 0.586 | 39.9 | 40.6 | 22.5 | 56.7 | 29.5 |
| **LitR** | 606 | 0.561 | 0.483 | 43.1 | 51.0 | 33.2 | 45.0 | 26.3 |
| **LmrA** | 567 | 0.556 | 0.463 | 46.7 | 57.7 | 42.3 | 40.2 | 23.1 |
| **QacR** | 567 | 0.641 | 0.576 | 39.2 | 39.7 | 23.3 | 54.5 | 28.4 |
| **SrpR** | 642 | 0.491 | 0.390 | 46.9 | 68.7 | 36.4 | 35.5 | 21.9 |
| **LacI** | 1083 | 0.531 | 0.462 | 56.1 | 63.4 | 43.2 | 61.8 | 48.0 |
| **YFP** | 720 | 0.867 | 0.865 | 61.0 | 57.1 | 30.8 | 95.0 | 23.7 |

Table 5: **Prediction metrics**.

| Gate | $R^2$ | MSE | RMSE | MAE |
|---|---|---|---|---|
| **LitRL1** | 0.86 | 0.00 | 0.04 | 0.03 |
| **HlyIIRH1** | -51.47 | 0.00 | 0.03 | 0.01 |
| **BetIE1** | -17.85 | 0.30 | 0.55 | 0.54 |
| **IcaRAI1** | -1.07 | 0.03 | 0.16 | 0.13 |
| **LmrAN1** | -0.03 | 0.01 | 0.12 | 0.12 |
| **QacRQ1** | 0.90 | 0.00 | 0.06 | 0.01 |
| **QacRQ2** | -0.73 | 0.14 | 0.37 | 0.36 |
| **SrpRS1** | 0.69 | 0.00 | 0.03 | 0.03 |
| **SrpRS2** | 0.91 | 0.00 | 0.03 | 0.03 |
| **SrpRS4** | 0.88 | 0.00 | 0.03 | 0.03 |

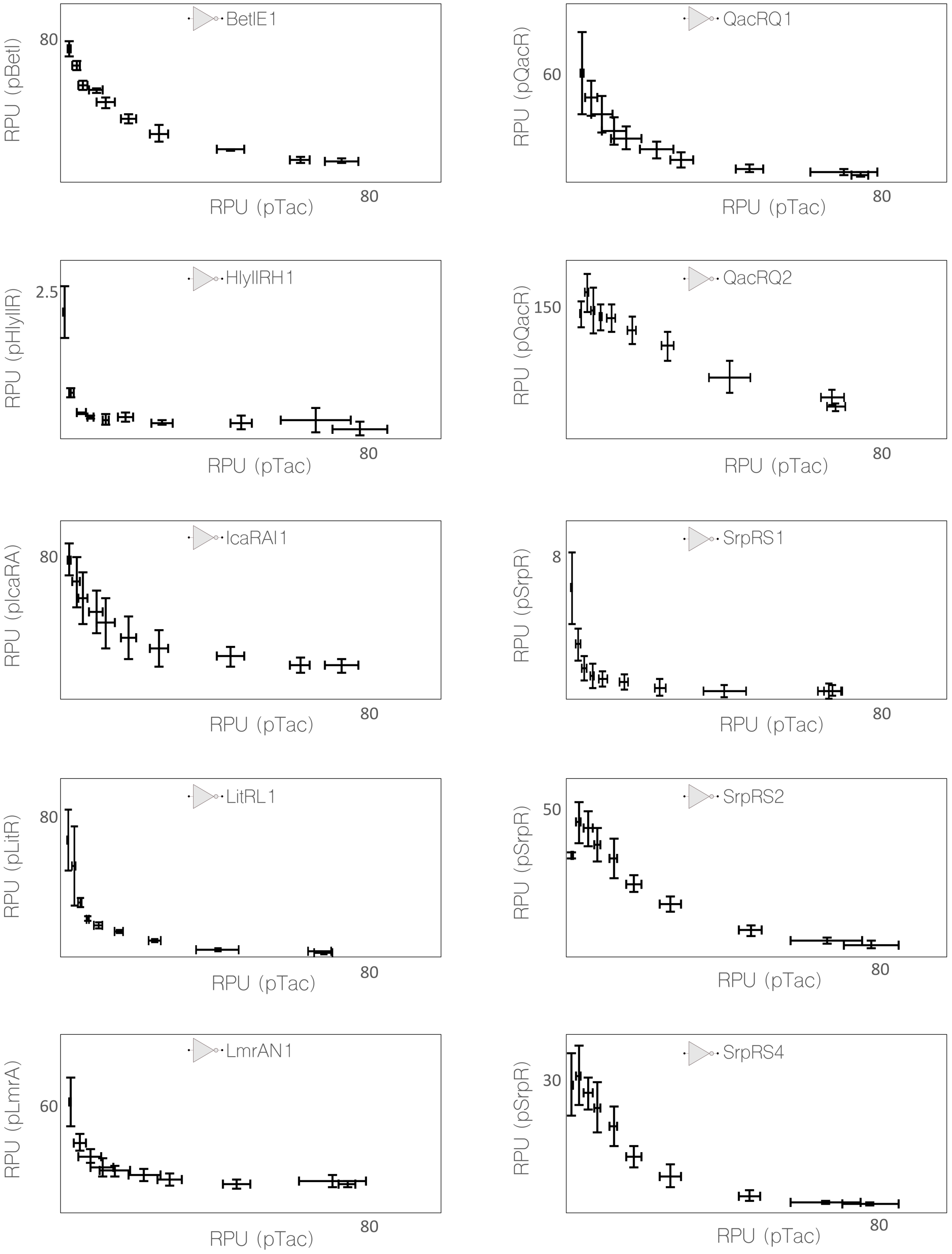


Figure 1: **Experimental characterization of the library of inverters**. The 10 genetic NOT gates from the benchmarking library were tested at increasing concentrations of IPTG ranging from 0 to 1000 µM. Input and output promoters activity was measured as relative units using flow cytometry fluorescence measurements for *Pseudomonas protegens* Pf-5. The NOT function is plotted as follows: the input promoter RPUs (pTac, x-axis) and the output promoters RPUs (pRx, y-axis) expressed as a percentage of the empty NOT gate fluorescence. Cells were incubated in M9 minimal medium supplemented with citrate and the corresponding IPTG concentration for 24 hours. Average and standard deviation were calculated from three independent biological replicates.

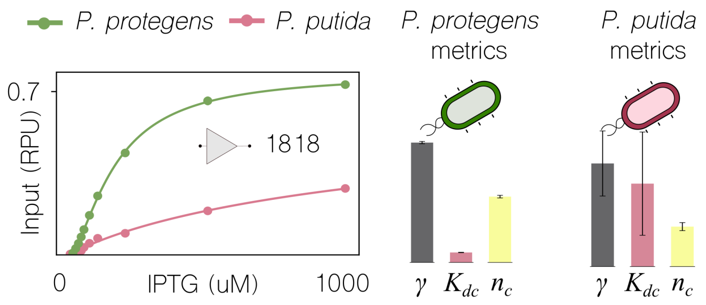


Figure 2: **Calibration of the IPTG sensor module LacI-pTac**. The input promoter (pTac) activity was characterized using input concentrations of IPTG ranging from 0 to 1000 µM. Output fluorescence was measured as relative units adjusting the standards to those of *P. protegens*, facilitating direct comparisons. The results were plotted against those of *P. putida*. In all plots, experimental data and model fit are depicted using dots and solid lines, respectively. Additionally, for each bacterial species, 3 testing metrics $\gamma$, $K_{dc}$ and $n_c$ were calculated and presented in bar plots. Parameter values except $K_{dc}$ (concentration arbitrary units) are dimensionless using intervals: 0-8(x$10^{-1}$) for $\gamma$, 0-2 (x$10^3$) for $K_{dc}$, and 0-3 for $n_c$. For *P. protegens*: $\gamma = 0.779 \pm 0.008, K_{dc} = 166 \pm 4, n_c = 1.52 \pm 0.03$. For *P. putida*: $\gamma = 0.7 \pm 0.2, K_{dc} = 1349 \pm 840, n_c = 0.90 \pm 0.09$.

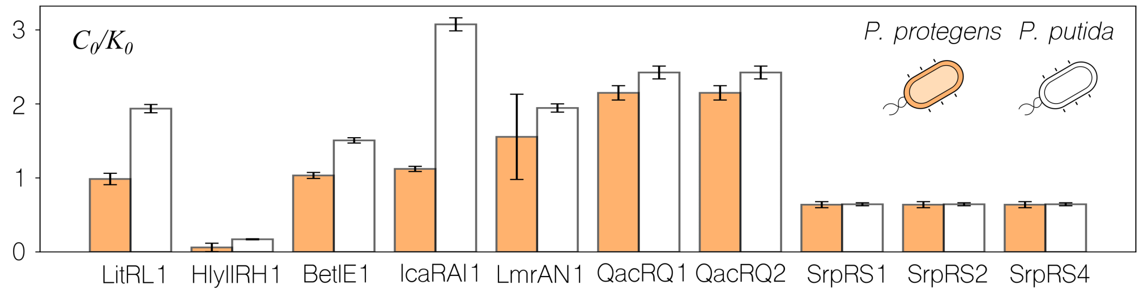


Figure 3: **Output promoter activity in *Pseudomonas protegens* Pf-5**. Characterization of the output promoters transcription rates (C0) normalized by the input promoter transcription rate (K0) in *P. protegens* Pf-5 (coloured). Depicted in white, the equivalent ratio for *Pseudomonas putida* KT2440. This ratio comes from $a/\gamma = C_0/K_0$.

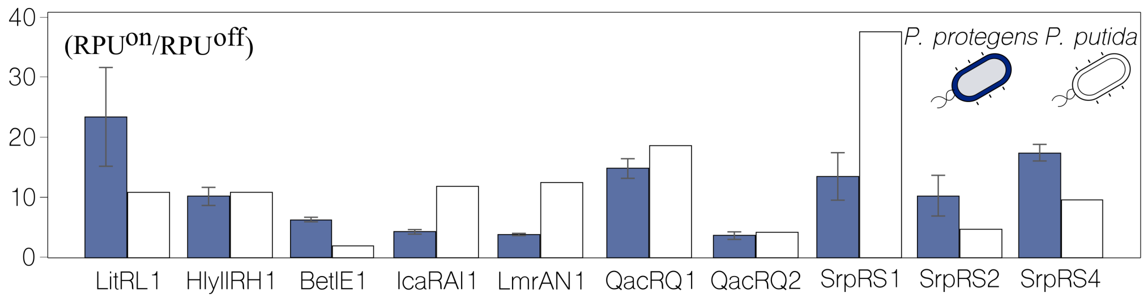


Figure 4: **Dynamic ranges of the benchmarking library of genetic NOT logic inverters**. The fold-change between the ON and OFF states of the 10 inverters was calculated by dividing the output RPUs at input 0 (low IPTG) by their equivalent at input 1 (high IPTG). The error bars for *Pseudomonas protegens* Pf-5 represent the standard deviation using the medians obtained by flow cytometry from at least 3 biological replicates. The dynamic ranges for *Pseudomonas putida* KT2440 were calculated utilizing previously published datasets [2].

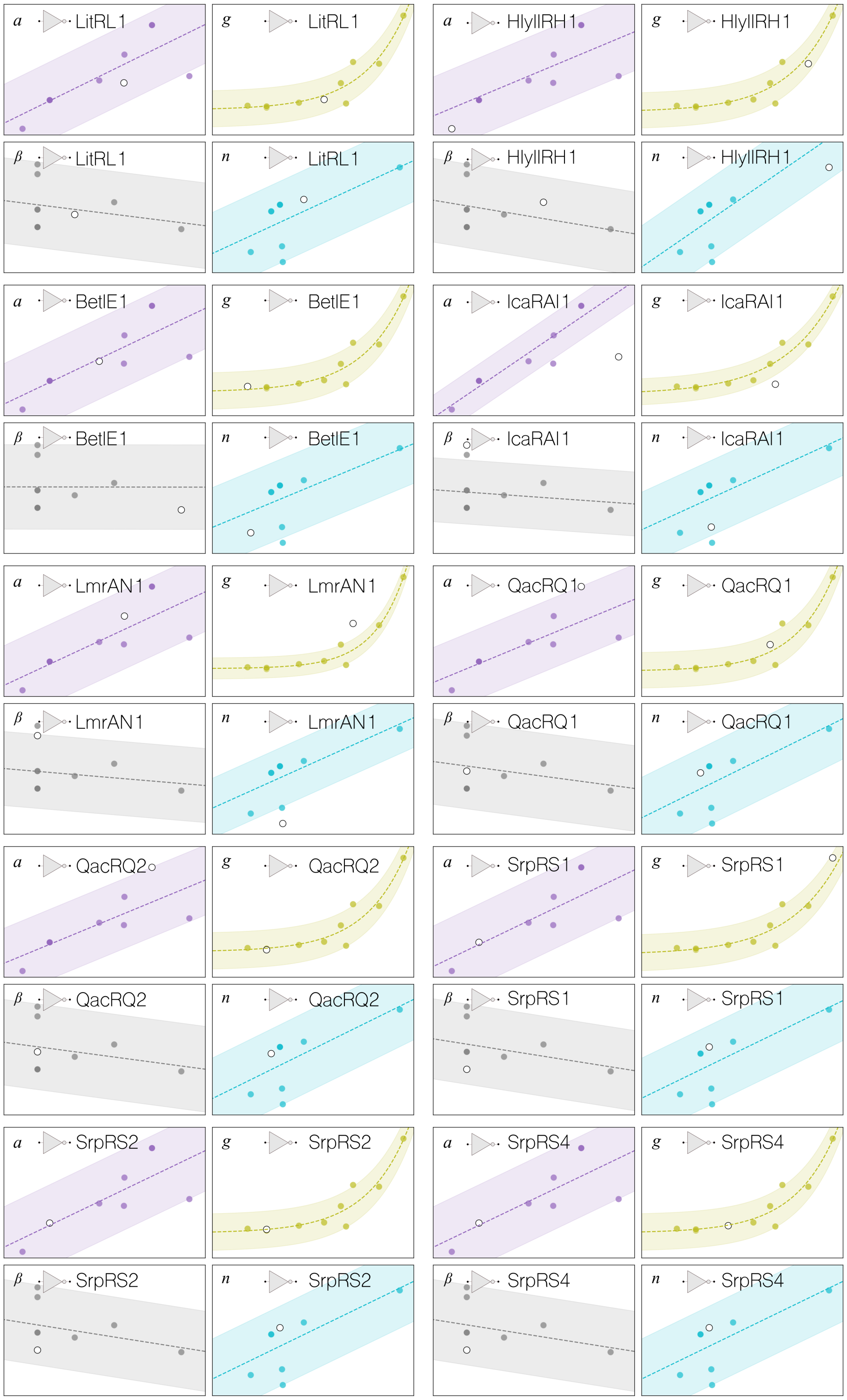


Figure 5: **Leave-One-Out Cross-Validation for the 10 inverters of the library**. Interspecies correlation of the inverters performance metrics between *P. putida* KT2440 (x-axis) and *P. protegens* Pf-5 (y-axis). The trajectory and the deviation for each parameter are represented as dashed lines and shadowed areas respectively. Solid dots mark the intersection for each inverter between both hosts - white dots correspond to the excluded gate, identified over each correlation.

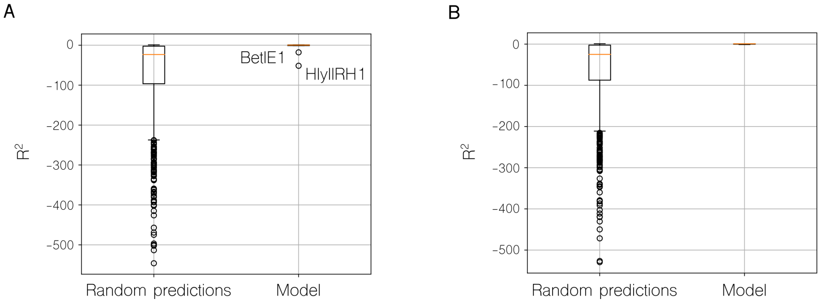


Figure 6: **Confidence assessment of randomized vs model predicted inverter dynamics**. **A**. Distribution of $R^2$ confidence assessment of random predictions against the model-based predictions of the 10 inverters of the library in *Pseudomonas protegens* Pf-5. Prediction outliers HlyIIRH1 and BetIE1 indicated. **B**. Distribution of $R^2$ confidence assessment of random predictions against the model-based predictions excluding the outliers. The probability that random predictions reach the confidence levels of the model-based predictions for the remaining 8 gates is 0.035 (below 0.05).
